# Posterior-calibrated multimodal motor states reveal longitudinal and imaging-associated heterogeneity in Parkinson’s disease

**DOI:** 10.64898/2026.06.12.732003

**Authors:** Harsh Milind Tirhekar, Priyanshi Yadav, Chandrajit Bajaj

## Abstract

Parkinson’s disease (PD) motor heterogeneity is commonly summarized by hard subtype labels, although clinical states vary longitudinally, severity can dominate unsupervised structure, and model uncertainty is rarely calibrated. We developed a posterior and refit-stability calibrated multimodal motor state framework that assigns probabilistic MDS-UPDRS-III motor states, aggregates them at the patient level, separates global burden from residual tremor-axial profile, and tests whether imaging can recover the resulting posterior distribution. In 29,366 aligned PPMI motor-posterior visits spanning 4,773 participant identifiers, patient-level state families were stable on average (modal-family fraction 0.925; 95% CI 0.921 - 0.930), but 25.5% of patients transitioned state over follow-up (95% CI 24.1 - 26.7%). PD-only cohort definitions produced smaller denominators and are reported as sensitivity cohorts with rerun calibration and imaging-posterior checks. Severity and covariates explained substantial motor-domain variance, especially bradykinesia (*R*^2^=0.850), but residual profile modeling retained five active components across total-severity, principal-component, leave-one-domain, non-target-burden, and clinical-only severity axes. Refit-stability calibration with 250 patient-blocked bootstrap refits showed high nominal posterior confidence (0.989) but lower empirical label consistency (0.849), quantifying overconfidence rather than hiding it. Patient-held-out temporal modeling predicted future axial burden (best XGBoost *R*^2^=0.605) and future state transition (XGBoost AUC=0.830; 95% CI 0.822 - 0.837). DaTSCAN plus FreeSurfer ROI features predicted patient-level soft motor posterior vectors (RF Jensen–Shannon divergence=0.209; 95% CI 0.199 - 0.220; macro-AUROC=0.692), while severity/demographic-adjusted imaging features further improved soft posterior recovery (Jensen–Shannon divergence=0.188). BioFIND transfer reproduced clinically meaningful endpoint gradients after state assignment in 225 external patients, supporting external face validity rather than definitive transportability. These results support PD motor phenotypic states as calibrated, dynamic, clinically interpretable profiles with convergent imaging associations, not as definitive biological subtypes.

## Introduction

Parkinson’s disease (PD) is not a single motor phenotype. Tremor, rigidity, bradykinesia, axial impairment, gait dysfunction, and bulbar symptoms vary between patients and over time [1, 2]. Conventional tremor-dominant and postural instability/gait difficulty groupings remain clinically useful, but they compress high-dimensional motor examinations into deterministic labels that can change with progression and assessment context [3, 6, 5, 4]. Large observational cohorts such as the Parkinson’s Progression Markers Initiative (PPMI) and BioFIND now make it possible to study this heterogeneity longitudinally, with clinical, dopaminergic imaging, structural MRI, and biomarker endpoints [7, 8, 9].

The central problem is not merely clustering. Prior work shows that subtype definitions can vary longitudinally and that severity levels are tightly coupled to MDS-UPDRS burden [6, 10, 16]. State-based and progression-subtype studies further show that PD trajectories can be overlapping, nonsequential, and multimodal rather than reducible to a single static label [17, 19, 20, 21]. A clinically useful motor phenotyping system should therefore answer four harder questions. First, does a model distinguish a patient-level phenotype family from a visit-level disease state? Second, does it separate global motor severity from residual profile structure, especially tremor versus axial burden? Third, are posterior probabilities reliable under patient-level refitting, or do they express mathematical confidence without empirical stability [26, 27, 28, 29]? Fourth, can multimodal imaging predict the posterior state distribution rather than only validate a hard label after the fact [11, 12, 13, 14, 22]?

A preliminary motor-only analysis established a computational base: a large Bayesian Gaussian mixture model (BGMM) sweep over 2,912 configurations, five macro-states, posterior assignment, DaTSCAN and FreeSurfer validation, and BioFIND transfer. The present work is a full journal-level expansion. We deliberately avoid the claim that five discovered clusters are “true PD subtypes.” Instead, we ask whether posterior-calibrated motor phenotypic states are stable at the patient level, meaningfully dynamic over visits, separable from severity, predictive of future motor change, recoverable from imaging, and externally associated with BioFIND clinical endpoints. This framing is aligned with recent data-driven progression studies and multimodal subtype literature while avoiding overinterpretation of unsupervised clusters [15, 16, 13].

Our framework, posterior-calibrated multimodal motor state modeling (PC-MSM), has seven components: patient-level posterior aggregation, visit-to-visit state dynamics, severity-deconfounded residual profiles, patient-blocked posterior calibration, future-state prediction, imaging-to-posterior soft-label learning, and external endpoint validation. The resulting model is positioned not as a replacement for clinical judgment, but as a calibrated computational phenotype layer for cohort stratification, trial enrichment, and imaging-aware PD progression analysis.

The study was organized around seven guardrails that directly address common weaknesses in unsupervised disease subtyping: patient-held-out evaluation rather than visit-independent leakage, patient-blocked bootstrap calibration rather than threshold-only posterior confidence, severity residualization and matched-state contrasts rather than unadjusted symptom burden, effect-size-first imaging validation rather than p-value counting, soft-label imaging prediction rather than hard-label post hoc testing, external BioFIND endpoint gradients rather than assignment confidence alone, and seed/baseline benchmarking rather than single-model reporting.

### Formal framework and terminology

For patient *i* at visit *v*, let *x_iv_* denote the MDS-UPDRS-III motor profile and *Z_iv_* ∈ {1*, . . ., K*} a latent motor state. The visit-level posterior is

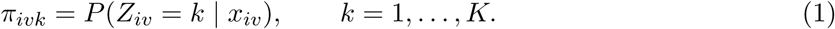

The patient-level phenotype-family posterior is the visit-averaged posterior

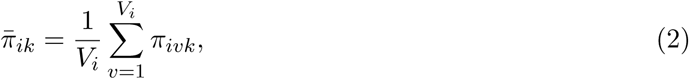

where *V_i_* is the number of observed visits. The modal patient family is arg max*_k_* 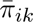, whereas visit-level states use arg max*_k_ π_ivk_*. This distinction is central: the model summarizes a patient’s dominant motor family without assuming that every visit belongs to an immutable subtype.

Because latent motor states have no biological gold-standard labels, calibration is defined operationally as refit stability, not as calibration to a true clinical class. After patient-blocked bootstrap refit *b* and Hungarian label alignment, empirical consistency for visit (*i, v*) is

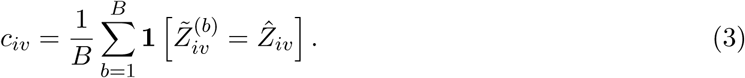

For confidence bin *B_m_*, stability calibration error is

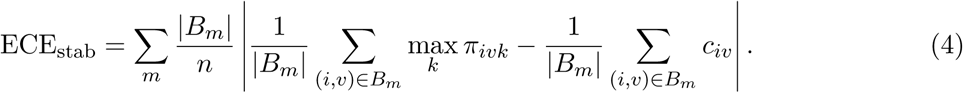

Thus, “posterior calibration” in this manuscript means posterior-vs-refit-stability reliability. Clinical calibration is reserved for supervised predictions such as next-visit transition and imaging-to-posterior prediction.

Severity-profile decomposition was defined by motor domain *d* as

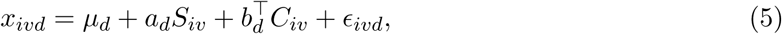

where *S_iv_* is global motor severity, *C_iv_* are covariates, and *ɛ_ivd_* is the severity-deconfounded residual profile. Imaging-to-posterior prediction estimates

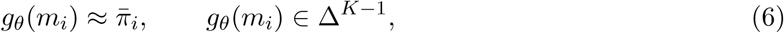

where *m_i_* are DaTSCAN and/or FreeSurfer ROI features. Evaluation used KL divergence, Jensen–Shannon divergence, Brier score, soft-label expected calibration error, and macro-AUROC, with reporting organized to align with modern guidance for clinical prediction models [30, 31].

## Results

### Analytic cohort audit and state language

The May 2026 pipeline realigned 29,366 complete MDS-UPDRS-III motor visits to the prior BGMM posterior assignment table, spanning 4,773 unique PPMI participant identifiers [7, 9]. This full aligned table was used for the computational framework. A separate cohort audit showed that stricter PPMI PD filters produce smaller denominators: 1,602 aligned patients under participant-status PD cohort filtering, 1,923 event-aligned patients under current primary clinical diagnosis PRIMDIAG == 1, and 1,123 patients under PD-cohort-and-enrolled filtering. Because multiple PPMI PD definitions are scientifically defensible, we report this inclusion audit explicitly rather than hardcoding a single retrospective denominator. All headline results below use the full aligned posterior table, while stricter PD-only sensitivity cohorts were rerun for posterior stability calibration and fused imaging-to-posterior prediction.

**Table 1:** Cohort-definition sensitivity. Modal-family and transition fractions come from saved patient-state outputs. Refit-stability and fused imaging Jensen–Shannon divergence were rerun inside each cohort definition using 250 patient-blocked BGMM refits and 5-fold patient-held-out imaging-to-posterior prediction.

| Cohort definition | Visits | Patients | Modal family | Transitioned | Residual $k$ | | Fused JSD |
| --- | --- | --- | --- | --- | --- | --- | --- |
| Full aligned table | 29,366 | 4,773 | 0.925 | 25.5% | 5 | 0.140 | 0.209 |
| PRIMDIAG == 1 | 19,032 | 1,923 | 0.830 | 58.0% | 5 | 0.123 | 0.246 |
| PD cohort | 16,899 | 1,602 | 0.820 | 61.6% | 5 | 0.152 | 0.246 |
| PD cohort + enrolled | 12,143 | 1,123 | 0.812 | 63.8% | 5 | 0.150 | 0.241 |

This audit prevents inflated certainty about the denominator and keeps the interpretation aligned with the available data: we study posterior-calibrated motor phenotypic states in the aligned PPMI motor-posterior table, with candidate PD-only filters documented as sensitivity cohorts.

### Patient-level posterior aggregation separates phenotype family from visit-level state

For each patient, visit-level posterior vectors were averaged to produce a patient-level posterior distribution. The modal patient state and modal phenotype family were then computed from this posterior trajectory. Across 4,773 patients, the mean modal-state fraction was 0.918 and the mean modal-family fraction was 0.925. Among 2,014 patients with at least five visits, longitudinal information was sufficiently dense for state-dynamic interpretation. Overall, 25.5% of patients changed motor state at least once, and mean dwell time was 3.35 visits.

These findings support the revised conceptual framing. The model does not define immutable biological subtypes. It identifies a stable-enough patient-level phenotype family while preserving visit-level motor-state dynamics. The state transition matrix provides a direct way to study disease-state evolution and mixed/posterior-boundary cases.

### Severity explains major motor-domain variance, but residual profile structure remains

A major risk in PD phenotyping is that clusters become discretized severity strata. We therefore residualized robust-scaled motor domains against total MDS-UPDRS-III severity, age, disease duration, sex, medication state, and levodopa equivalent daily dose (LEDD) when available. Severity and covariates explained substantial domain variance: bradykinesia *R*^2^=0.850, rigidity *R*^2^=0.696, bulbar *R*^2^=0.674, axial *R*^2^=0.448, and tremor *R*^2^=0.423. This confirms that severity is not a nuisance detail; it is a dominant axis that must be modeled.

After residualization, a BGMM retained five active components. Agreement between raw and total-severity residual labels was modest (adjusted Rand index=0.228, normalized mutual information=0.147), indicating that raw symptom burden and residual motor profile capture different axes. The finding was not unique to the total-score severity axis: five active residual components were also retained when severity was represented by the first motor-domain principal component (adjusted Rand index=0.337), leave-one-domain total burden (adjusted Rand index=0.394), non-target-domain burden (adjusted Rand index=0.394), or clinical/covariate-only burden (adjusted Rand index=0.369). A matched severe-tremor versus severe-axial comparison produced 48 patient pairs after matching on severity, age, and disease duration. The key figure plots global motor severity against tremor-minus-axial profile contrast. Vertical separation at similar severity supports profile structure beyond total burden and reduces the part-whole artifact concern.

**Figure 1:**
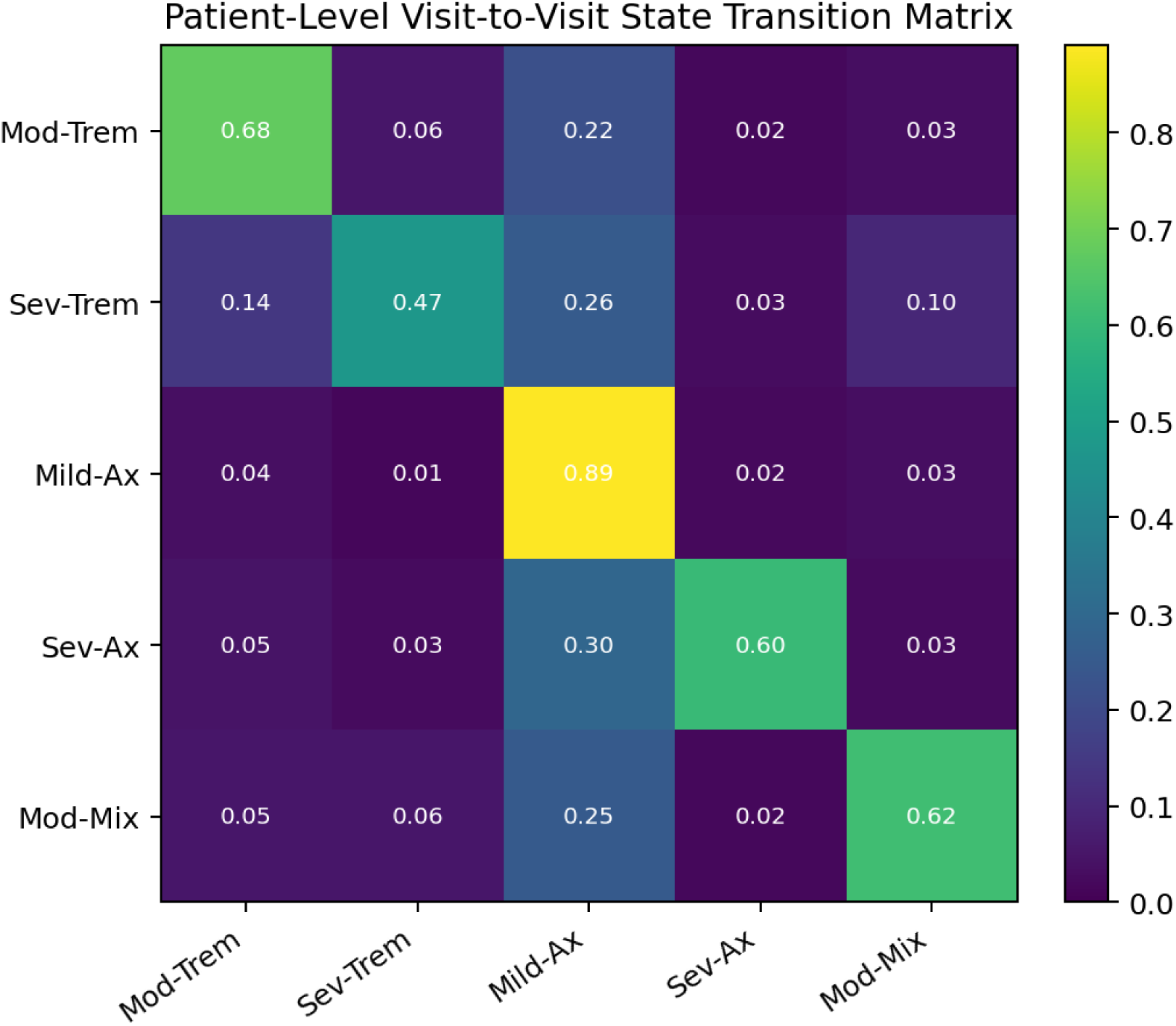
Patient-level motor-state dynamics. Visit-to-visit transition probabilities across the five posterior motor states, estimated from longitudinal PPMI motor visits. Rows indicate current state and columns indicate next observed state.

### Refit-stability calibration quantifies uncertainty rather than relying on arbitrary thresholds

The original posterior triage concept was clinically useful, but threshold-only language can be misleading if posterior probabilities are overconfident [26]. We performed 250 patient-blocked bootstrap refits of the selected BGMM configuration, following the principle that cluster stability should be assessed under perturbation rather than assumed from a single fit [32, 33, 34]. For each refit, labels were aligned to the original state space using the Hungarian algorithm, and each visit’s empirical consistency was defined as the proportion of bootstrap refits preserving its aligned state.

**Figure 2:**
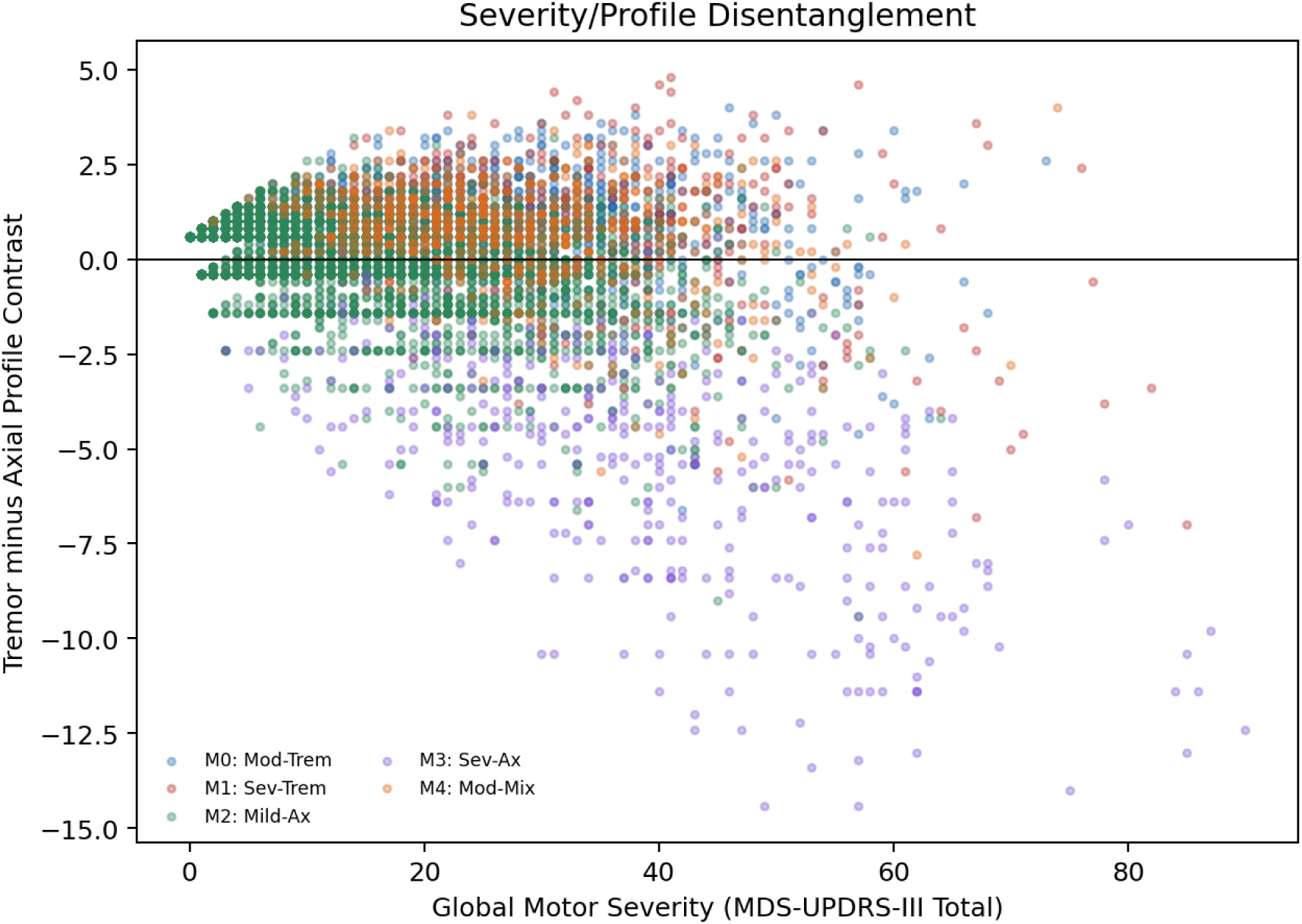
Severity/profile disentanglement. Global MDS-UPDRS-III severity is shown against tremor-minus-axial profile contrast. Separation along the vertical axis at similar severity supports residual profile information beyond total motor burden.

The model’s mean max-posterior confidence was 0.989, but mean bootstrap label consistency was 0.849. The full-table was 0.140 and the Brier-like inconsistency score was 0.084 (95% CI 0.080 - 0.090). PD-only sensitivity cohorts showed the same qualitative pattern: =0.123 for PRIMDIAG == 1, =0.152 for the PD cohort, and =0.150 for the PD-cohort-and-enrolled subset. Thus, the model is highly decisive mathematically, but empirical refit stability is lower. This is not a failure; it is exactly the reason to report refit-calibrated uncertainty. Posterior entropy and posterior gap become continuous uncertainty measures rather than coarse “textbook” or “ambiguous” bins.

### Posterior uncertainty and motor profile predict future state change

To replace causal Granger language, we trained patient-held-out temporal models to predict future axial burden and future state transition. The longitudinal table contained 24,593 next-visit prediction rows from 3,775 patients. Strong tabular baselines were evaluated in addition to random forests. Using current motor domains, severity, posterior probabilities, posterior uncertainty, covariates, and visit gap, XGBoost predicted next-visit axial burden with *R*^2^=0.605, RMSE=1.148, and MAE=0.683; LightGBM achieved *R*^2^=0.599; random forest achieved *R*^2^=0.593; and ridge regression achieved *R*^2^=0.586. The autoregressive baseline using current axial burden alone achieved *R*^2^=0.505 and RMSE=1.285. A patient-clustered Gaussian GEE baseline achieved in-sample *R*^2^=0.532.

Future state transition occurred in 19.0% of prediction rows. The all-feature XGBoost classifier achieved AUC=0.830 (95% CI 0.822 - 0.837), random forest achieved AUC=0.827 (95% CI 0.819 - 0.835), LightGBM achieved AUC=0.824 (95% CI 0.816 - 0.832), and logistic regression achieved AUC=0.781 (95% CI 0.771 - 0.791). Posterior-vector-only models remained predictive (XGBoost AUC=0.793), and entropy/gap-only models retained signal (LightGBM AUC=0.765; XGBoost AUC=0.762). A patient-clustered binomial GEE achieved AUC=0.698, and a Cox transition hazard model had concordance 0.620; higher severity and posterior entropy were associated with greater transition hazard. Time-gap-stratified transition matrices and a categorical HMM provided additional generative trajectory summaries.

**Figure 3:**
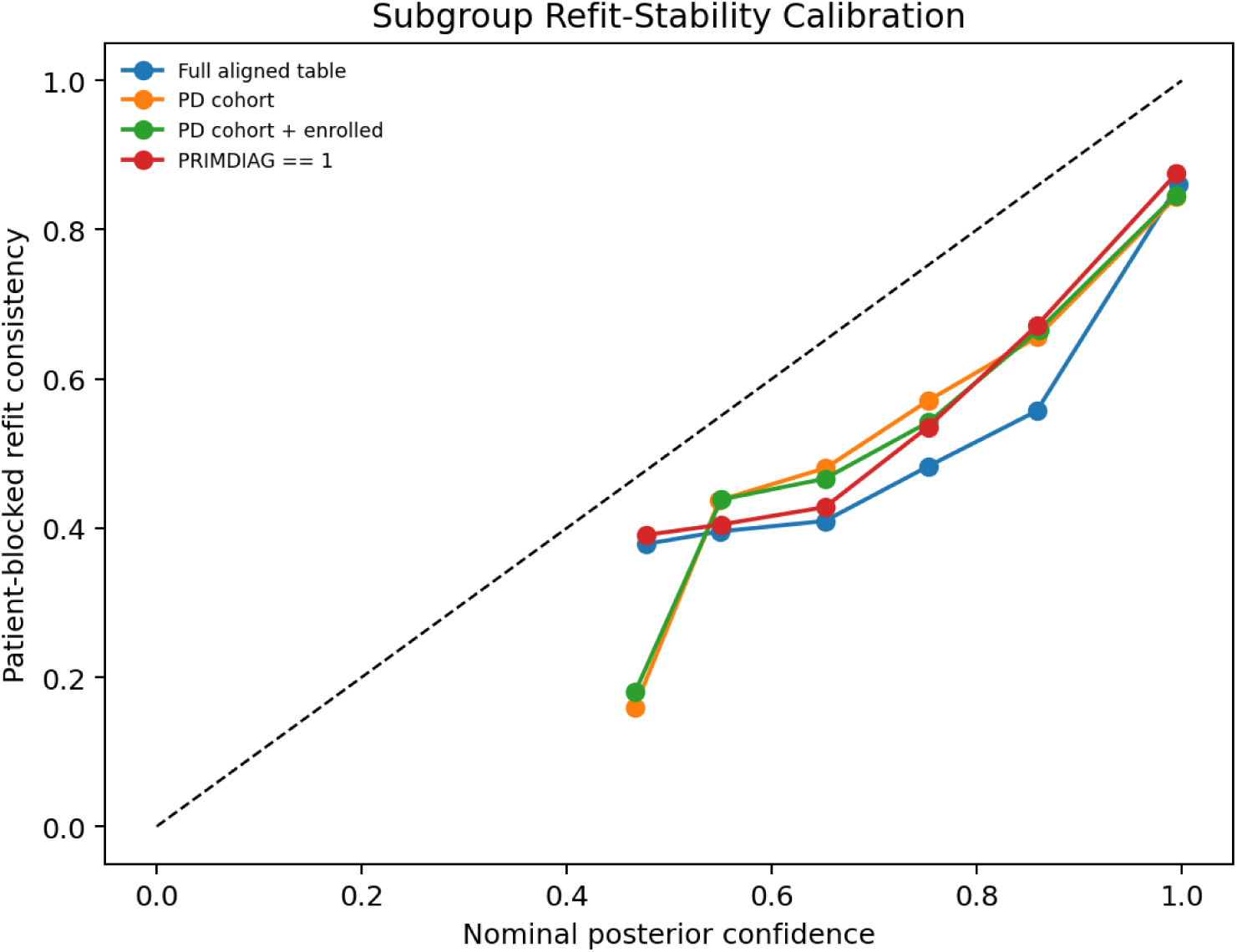
Cohort-specific refit-stability calibration. Nominal posterior confidence is compared with patient-blocked bootstrap consistency across the full aligned table and stricter PD-only cohort definitions.

These results give the posterior model clinical meaning: posterior uncertainty is associated with, and predicts, next-visit transition under patient-held-out validation. This is a predictive association rather than a causal claim.

### Imaging predicts soft motor posterior distributions

A key journal-level contribution is the shift from imaging as downstream validation to imaging as a posterior prediction problem. Prior neuroimaging literature supports dopaminergic, network-level, and structural differences between motor subtypes, but findings are heterogeneous and often modality-specific [11, 12, 13, 14, 22]. We trained RF, XGBoost, and LightGBM soft-label models to predict patient-level motor posterior vectors from DaTSCAN ROI features, FreeSurfer MRI ROI features, or fused DaTSCAN+MRI ROI features under 5-fold patient-held-out cross-validation.

In the full aligned imaging subset, fused DaTSCAN+MRI features in 1,041 patients produced the strongest pure imaging model with RF: KL=0.567, Jensen–Shannon divergence=0.209 (95% CI 0.199 - 0.220), Brier=0.251, soft-label expected calibration error=0.034, and macro-AUROC=0.692. XGBoost had similar hard-state discrimination (macro-AUROC=0.692) but weaker soft-label divergence (Jensen–Shannon divergence=0.215), and LightGBM achieved Jensen–Shannon divergence=0.220. Stricter PD-only sensitivity cohorts remained predictive but weaker: RF fused Jensen–Shannon divergence=0.246 for PRIMDIAG == 1, 0.246 for the PD cohort, and 0.241 for PD-cohort-and-enrolled. Feature-group ablations showed that DaTSCAN-only features (Jensen–Shannon divergence=0.213) carried more signal than MRI-only features (Jensen–Shannon divergence=0.266), while severity/demographic-plus-imaging features improved soft posterior recovery (Jensen–Shannon divergence=0.188; expected calibration error=0.015). Imaging residualized for severity retained signal (Jensen–Shannon divergence=0.206; macro-AUROC=0.739).

**Figure 4:**
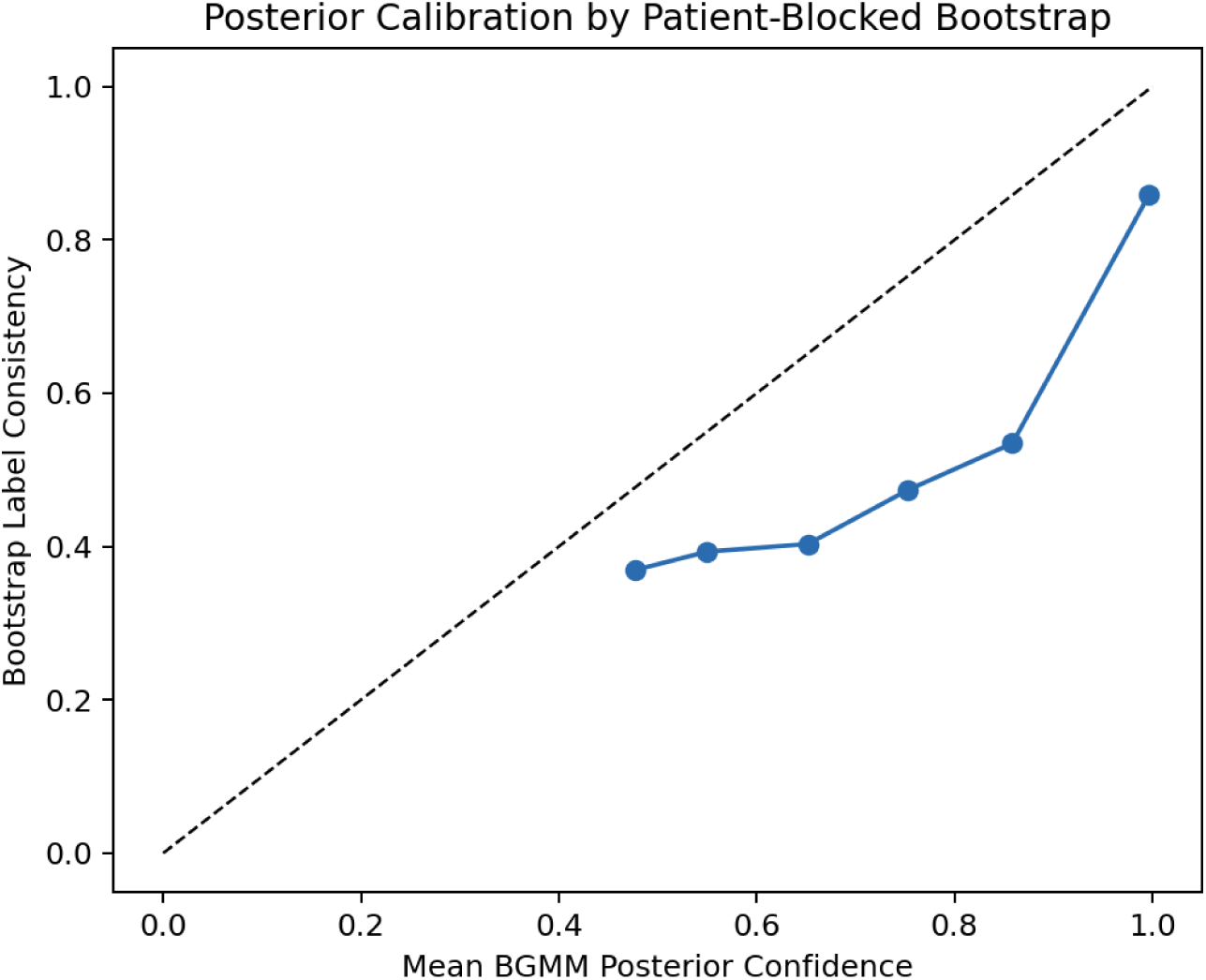
Refit-stability calibration. Mean BGMM posterior confidence is compared against empirical label consistency from 250 patient-blocked bootstrap refits. The calibration target is refit stability, not a biological ground-truth state label.

Feature attribution was biologically plausible and was evaluated using held-out permutation importance and SHAP, not only impurity-based importance [35, 25]. Permuting mean putamen SBR produced the largest held-out JSD increase (ΔJSD=0.036), followed by caudate asymmetry (0.013), anterior putamen asymmetry (0.012), caudate-putamen ratio (0.011), putamen asymmetry (0.008), and left/right putamen SBR. SHAP summaries agreed: mean putamen SBR was the leading predictor, followed by caudate asymmetry, anterior putamen asymmetry, left putamen SBR, putamen asymmetry, and caudate-putamen ratio, with selected structural MRI contributions from hippocampus, accumbens, third ventricle, thalamus, caudate, and related subcortical regions.

This supports an imaging-associated motor-state profile, not an imaging-defined ground truth. The strongest claim is that ROI-level imaging can recover clinically meaningful posterior information, especially when evaluated with soft-label calibration metrics.

**Figure 5:**
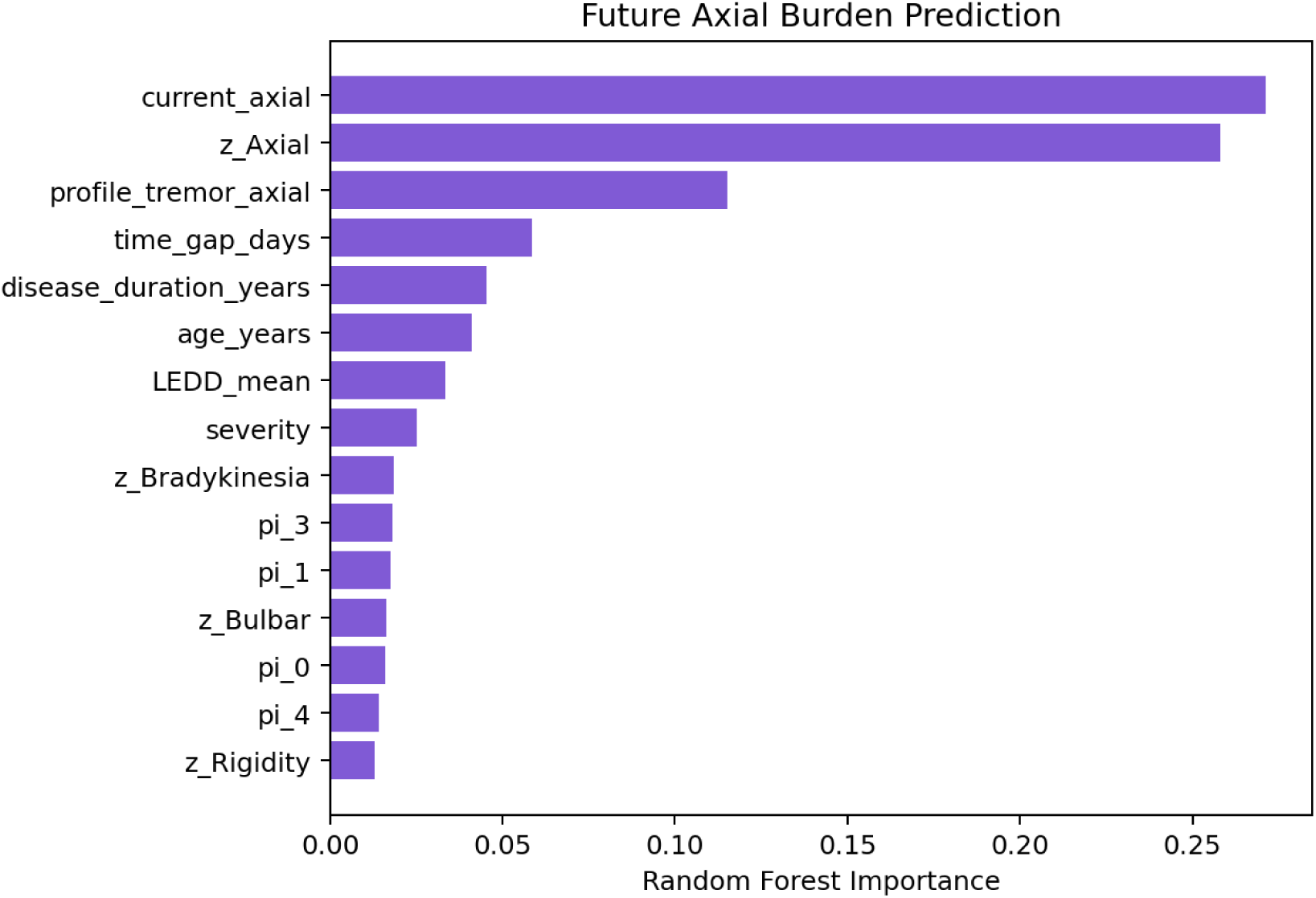
Longitudinal predictive features. Random-forest feature importance for predicting next-visit axial burden under patient-held-out cross-validation.

### Dopaminergic asymmetry dominates effect-size-first imaging validation

We next evaluated imaging endpoints using effect sizes, not p-values alone. Covariate-adjusted Δ*R*^2^ quantified how much patient-level motor state improved imaging prediction beyond severity, age, and disease duration. DaTSCAN asymmetry produced the strongest incremental effects: putamen asymmetry Δ*R*^2^ = 0.200 and *η*^2^ = 0.206; caudate asymmetry Δ*R*^2^ = 0.175 and *η*^2^ = 0.193. Mean putamen and caudate SBR also showed state-associated signal, but much smaller incremental effects after covariates.

Structural MRI effects were smaller, as expected for an early/progression cohort. Left thalamus, right thalamus, third ventricle, and hippocampal normalized volumes showed modest Δ*R*^2^ values. These results support a conservative interpretation: dopaminergic asymmetry is the strongest imaging-associated signal, while MRI contributes modest complementary information.

### BioFIND transfer shows external clinical endpoint gradients

Because BioFIND has no ground-truth motor-state labels, external validation was framed as endpoint validation after transfer rather than as proof of a subtype taxonomy. We fit the PPMI transfer BGMM/scaler and assigned BioFIND post-dose MDS-UPDRS-III visits. The external table contained 344 BioFIND motor visits from 225 patients. Mean BioFIND max posterior was 0.958 and mean entropy was 0.108. The PPMI-versus-BioFIND patient-state distribution Jensen–Shannon divergence was 0.069, indicating modest distribution shift.

Across clinical endpoints, assigned state was strongly associated with BioFIND motor severity (bootstrap mean *η*^2^ = 0.628, 95% CI 0.537 - 0.712), Hoehn and Yahr stage (*η*^2^ = 0.568, 95% CI 0.458 - 0.679), axial domain burden (*η*^2^ = 0.550, 95% CI 0.388 - 0.693), tremor (*η*^2^ = 0.541, 95% CI 0.436 - 0.630), rigidity (*η*^2^ = 0.539, 95% CI 0.443 - 0.635), bulbar burden (*η*^2^ = 0.517, 95% CI 0.408 - 0.634), and bradykinesia (*η*^2^ = 0.513, 95% CI 0.396 - 0.621). The tremor-minus-axial profile also showed a robust endpoint gradient (*η*^2^ = 0.370, 95% CI 0.206 - 0.537), supporting profile information beyond a single global burden axis in the external cohort. MDS-UPDRS Part II showed a smaller but nonzero endpoint gradient (*η*^2^ = 0.230, 95% CI 0.117 - 0.352), and MoCA effects were small.

**Figure 6:**
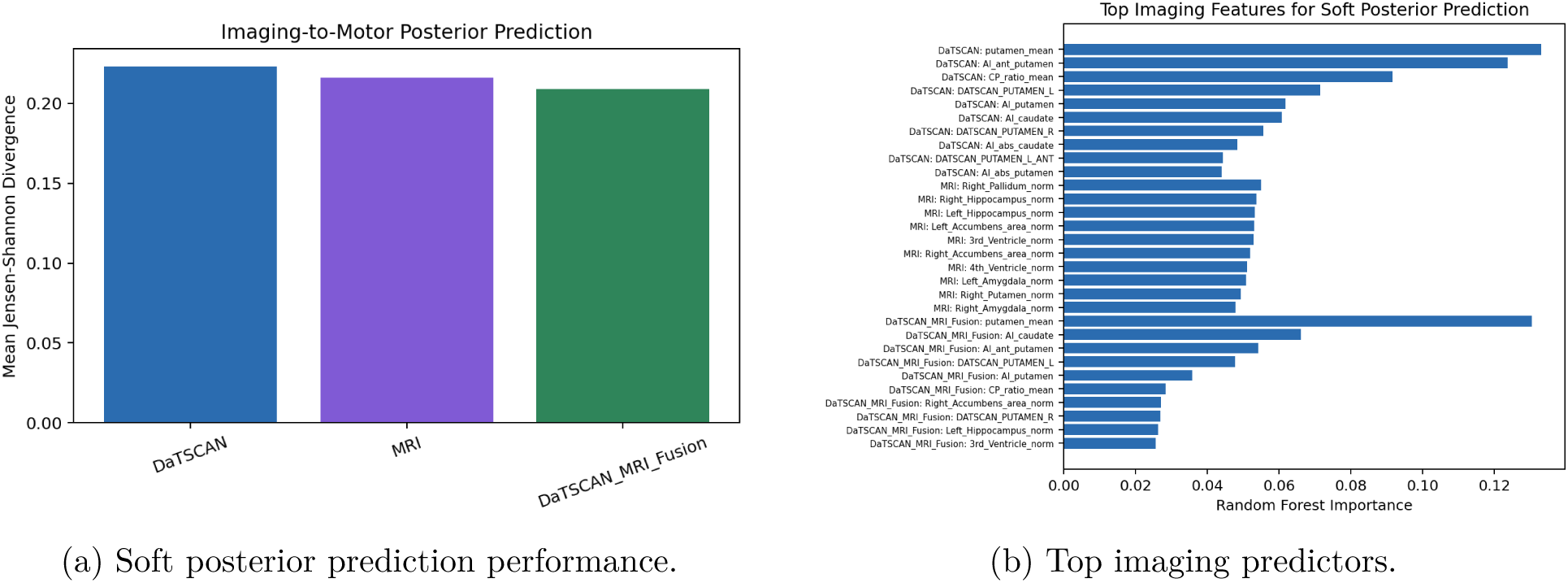
Imaging-to-posterior prediction. Patient-held-out soft-label prediction performance for DaTSCAN, MRI, and fused DaTSCAN+MRI models, together with random-forest feature attribution.

This analysis reframes BioFIND transfer as external clinical face validity and distribution-shift assessment.

### Imaging predicts motor posteriors but does not define the same latent taxonomy

We tested whether imaging-only unsupervised latent structure independently recovers the motor posterior states. It does not. Motor-only BGMM agreement with the prior motor state labels was ARI=0.451 and NMI=0.268. DaTSCAN-only latent mixtures had ARI=0.005 and NMI=0.121. MRI-only mixtures had ARI=0.023 and NMI=0.007. Inner fused motor+DaTSCAN+MRI mixtures had ARI=0.015 and NMI=0.116, while missing-aware multimodal mixtures had ARI=0.116 and NMI=0.091.

This negative result is important. Supervised imaging-to-posterior models can predict soft motor posteriors with moderate discrimination and good calibration, but unsupervised imaging clusters do not reproduce the same motor-derived taxonomy. The interpretation is therefore that motor states and imaging structure are related but not interchangeable.

### Baselines and practical runtime

The baseline benchmark used 75 seeds per stochastic method. BGMM was perfectly seed-stable in this configuration (pairwise ARI=1.000; pairwise NMI=1.000). K-means was also highly seed-stable (pairwise ARI=0.997; pairwise NMI=0.994), but less aligned to the original posterior states (ARI=0.234). Standard GMM was less stable (pairwise ARI=0.751; pairwise NMI=0.784). Agglomerative clustering had ARI=0.259 versus the original states.

The full May 2026 pipeline used 120 workers, 250 bootstrap refits, 5 cross-validation folds, 500 random-forest estimators, and 75 baseline seeds per method, completing in 119.48 seconds on the 144-core Vista node. This distinguishes the offline discovery sweep from practical inference and reproducible analysis iteration.

**Figure 7:**
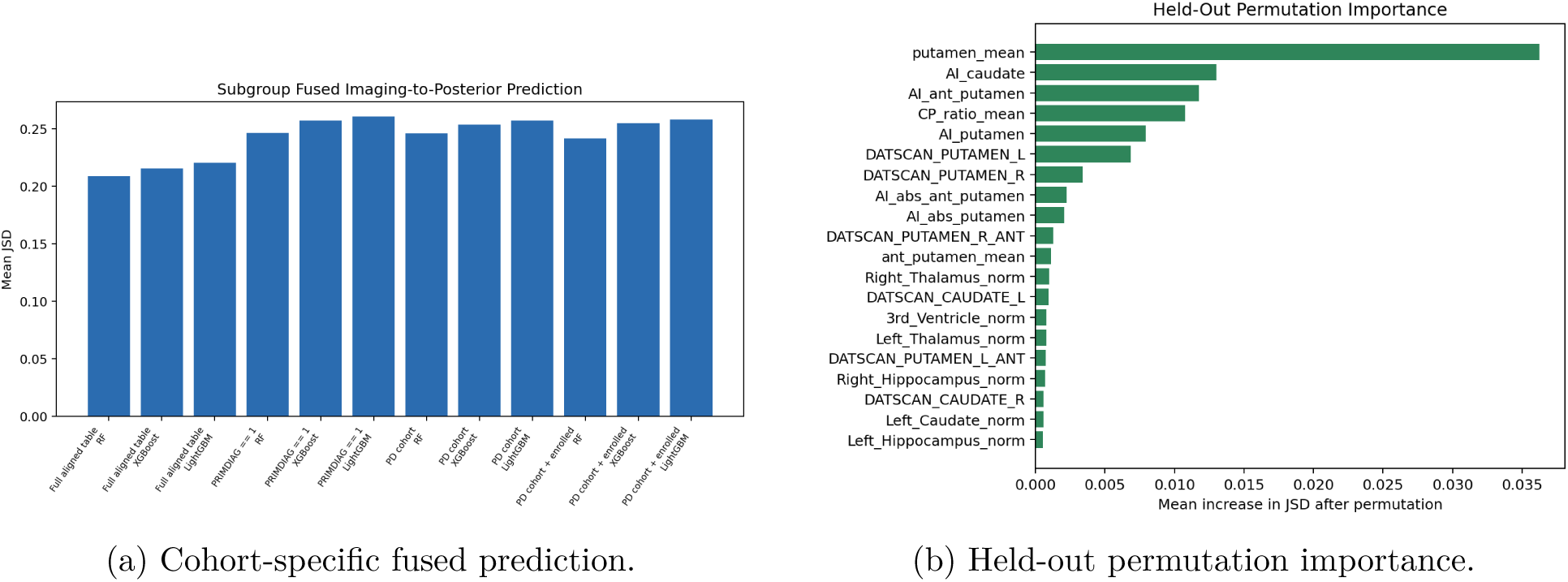
Professor-review imaging expansion. Subgroup fused imaging-to-posterior performance and foldwise permutation importance confirm that dopaminergic putamen burden and asymmetry dominate posterior recovery.

## Discussion

This study advances PD motor phenotyping in three ways. First, it changes the unit of interpretation from hard visit-level clusters to posterior-calibrated patient-level motor-state trajectories. Second, it makes severity a modeled axis rather than a confound hidden inside clustering. Third, it introduces a multimodal soft-label prediction framework in which DaTSCAN and structural MRI features predict the posterior distribution of motor states rather than merely testing hard-label differences after discovery.

The strongest clinical message is that hard subtype labels are the wrong unit for PD motor heterogeneity. A better unit is a calibrated posterior trajectory: stable enough to summarize a patient’s dominant motor family, dynamic enough to track transitions, severity-aware enough to avoid discretizing burden, and soft enough to be predicted from imaging and externally validated through endpoint gradients. Most patients have a dominant state family, but a substantial minority transition. Posterior uncertainty is associated with future transition risk, and future axial burden is predictable better than an autoregressive baseline when motor domains, posterior probabilities, and covariates are used together. This suggests a practical application in trial stratification: patients with high posterior uncertainty or mixed motor profiles may be enriched for near-term state change. The imaging results are also nuanced. Dopaminergic asymmetry, especially putamen and caudate asymmetry, shows the strongest incremental association with motor states. Fused DaTSCAN+MRI models produce the best calibrated soft posterior prediction. Yet imaging-only unsupervised clusters do not reproduce motor-state labels. Imaging therefore provides convergent and predictive evidence, not a ground-truth taxonomy.

External BioFIND analysis strengthens the framework by moving beyond decisive assignment. Endpoint gradients in severity, axial burden, Hoehn and Yahr stage, motor domains, and MDS-UPDRS Part II show clinical face validity, while distribution-shift metrics keep the interpretation honest. BioFIND validates endpoint gradients after transfer, not general-purpose transportability of the posterior state model.

**Figure 8:**
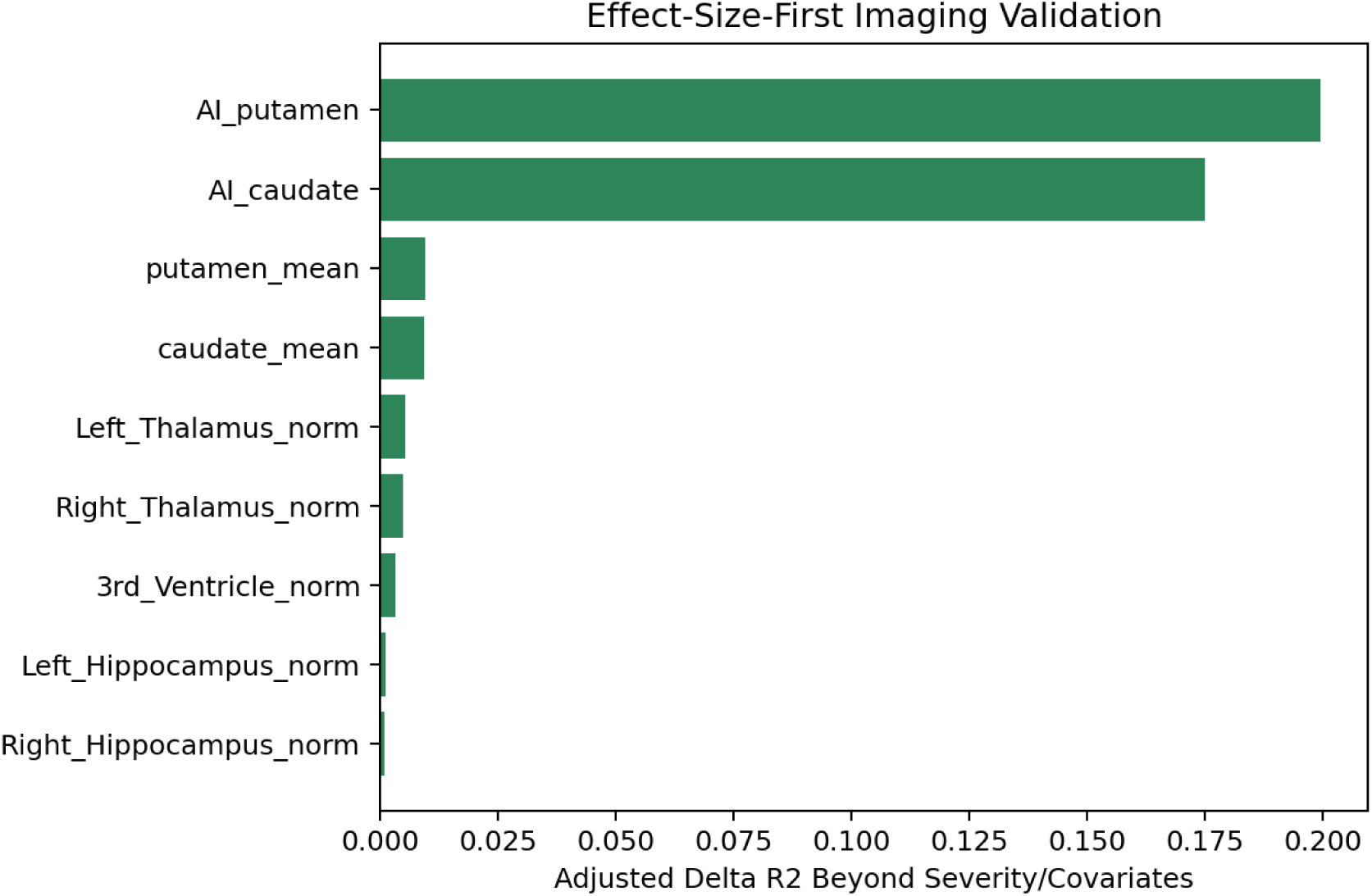
Effect-size-first imaging validation. Covariate-adjusted Δ*R*^2^ for imaging endpoints, emphasizing dopaminergic asymmetry as the strongest incremental imaging signal.

Several limitations remain. The current aligned table includes 4,773 PPMI participant identifiers, while stricter PD-only definitions yield smaller denominators and lower modal-family fractions; we therefore report PD-only sensitivity calibration and fused imaging prediction, but a final target-journal submission should prespecify one primary clinical cohort definition. Second, the frozen BGMM state space was discovered before the present downstream validation. A fully nested train-patient BGMM discovery, test-patient posterior assignment, and downstream validation loop remains the strongest leakage sensitivity. Third, the transition analyses now include time-gap-stratified matrices, Cox transition hazard, clustered GEE, and a categorical HMM, but they are still not a full irregular-time continuous-time Markov or semi-Markov disease-state model with transition intensities and dwell times in months or years [17, 18]. Fourth, the imaging-to-posterior models use ROI-level DaTSCAN and FreeSurfer features. This is interpretable and statistically tractable, but raw-image CNN or transformer models could test whether voxel-level information improves posterior prediction. Fifth, BioFIND endpoint validation is limited by available sample size and modalities. Finally, the framework is retrospective and requires prospective validation before clinical deployment.

In summary, posterior-calibrated multimodal motor state modeling provides a robust framework for PD motor heterogeneity. It yields patient-level state families, visit-level dynamics, severity-deconfounded profiles, calibrated posterior uncertainty, future-state prediction, imaging-posterior recovery, and external endpoint validation. The result is not a claim that PD has exactly five true biological subtypes. It is a clinically useful, uncertainty-aware model of motor phenotypic states that can support translational cohort stratification and mechanistic hypothesis generation.

**Figure 9:**
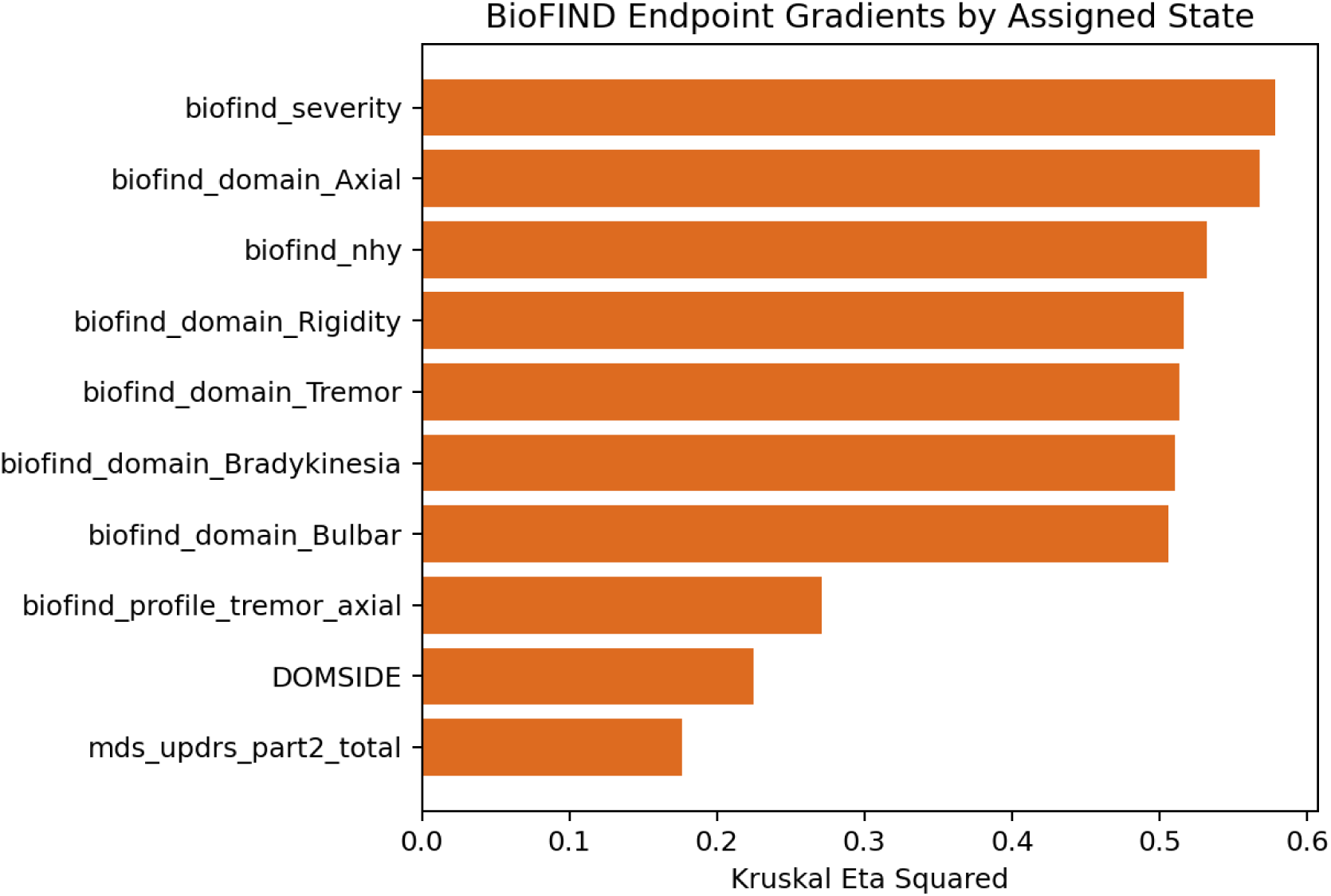
BioFIND endpoint gradients. External endpoint effect sizes by assigned BioFIND state, showing clinical face validity and severity-adjusted profile structure.

## Methods

### Data sources

The primary analysis used PPMI MDS-UPDRS Part III motor examination data and prior BGMM posterior assignments generated by the preliminary motor-state discovery analysis [2, 7, 9]. The motor table was filtered to complete NP3 item rows and aligned row-wise to bgmm_posterior_assignments_20260219_101 yielding 29,366 motor visits from 4,773 unique participant identifiers. Demographics, diagnosis history, medication fields, DaTSCAN SBR features, and FreeSurfer ASEG ROI features were loaded from the PPMI full data export available in the local project. External validation used BioFIND post-dose MDS-UPDRS Part III, MDS-UPDRS Parts I and II, MoCA, Hoehn and Yahr, and PD feature files when available [36, 37].

### Cohort inclusion audit

Because PPMI contains multiple participant categories and diagnosis tables, we audited multiple candidate PD filters without hardcoding an expected denominator. Participant-status filters included Parkinson’s Disease cohort, enrolled status, and their intersection. Diagnostic filters included current primary clinical diagnosis PRIMDIAG == 1, clinical diagnosis NEWDIAG == 1, and archived primary diagnosis PRIMDIAG == 1. For tables with EVENT_ID, both patient-level and event-aligned counts were reported. A follow-up sensitivity table was generated from saved per-patient and per-visit outputs for metrics that do not require rerunning the fitted models.

**Figure 10:**
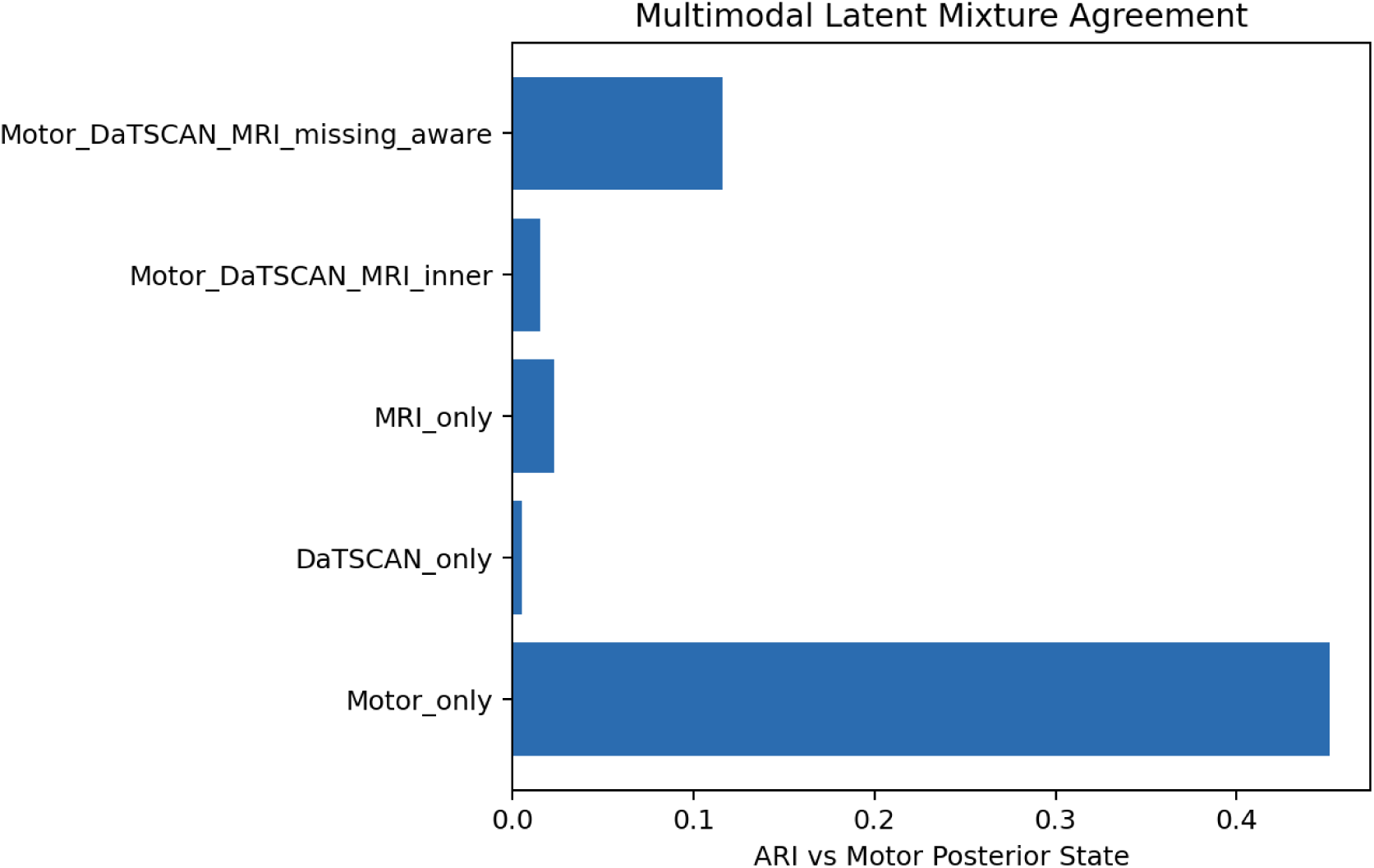
Multimodal latent mixture ablation. Agreement between motor posterior states and unsupervised latent mixtures from motor-only, imaging-only, fused, and missing-aware multimodal views.

### Motor domains

MDS-UPDRS-III items were grouped into tremor, rigidity, bradykinesia, axial, and bulbar domains. Domain scores were computed as item means over available domain items. Domain features were robust-scaled before residual modeling and patient-level summaries. Total motor severity used NP3TOT when present or the sum of NP3 item scores otherwise.

### Prior BGMM posterior model

The starting posterior assignments came from a previous BGMM discovery sweep across 2,912 configurations. BGMMs provide posterior probabilities and shrinkage over mixture components under a variational Bayesian formulation [23, 24]. The selected model provided five posterior state probabilities per motor visit. The present manuscript does not interpret these as definitive biological subtypes; they are used as probabilistic motor-state assignments for calibration, longitudinal modeling, imaging prediction, and external endpoint validation.

### Patient-level posterior aggregation and state dynamics

For each patient, visit-level posterior vectors were averaged and normalized to obtain a patient-level posterior. Modal state, modal family, first state, last state, state transition, posterior entropy, posterior gap, and mean max posterior were computed. Visit-to-visit transition counts were accumulated in a 5-by-5 transition matrix and normalized by origin state. Dwell time was computed as the length of consecutive same-state runs.

**Table 2:** Summary of headline verified analyses. All values come from the May 2026 full run in Updated_MICCAI_May2026.

| Analysis | Verified result |
| --- | --- |
| Patient-level stability | Mean modal-state fraction 0.918; mean modal-family fraction 0.925 (95% CI 0.921 - 0.930); 25.5% ever transitioned (95% CI 24.1 - 26.7%); mean dwell time 3.35 visits. |
| Severity deconfounding | Residual BGMM effective $k = 5$ under total, PC1, leave-one-domain, non-target-burden, and clinical-only severity axes; total-severity raw-vs-residual ARI=0.228. |
| Refit-stability calibration | 250 patient-blocked refits; =0.140; mean posterior confidence 0.989; mean bootstrap consistency 0.849; Brier-like score 0.084 (95% CI 0.080 - 0.090). |
| Temporal prediction | Future axial burden best XGBoost $R^2=0.605$; transition XGBoost AUC=0.830 (95% CI 0.822 - 0.837), RF AUC=0.827 (95% CI 0.819 - 0.835), Cox concordance=0.620. |
| Imaging posterior prediction | Fused DaTSCAN+MRI RF: JSD=0.209 (95% CI 0.199 - 0.220), macro-AUROC=0.692, soft-label ECE=0.034; severity/demographic+imaging JSD=0.188. |
| BioFIND transfer | 344 visits / 225 patients; PPMI-vs-BioFIND distribution JSD=0.069; BioFIND severity $\eta^2 = 0.628$ (95% CI 0.537 - 0.712), axial $\eta^2 = 0.550$ (95% CI 0.388 - 0.693). |
| Baselines | BGMM pairwise ARI=1.000 across 75 seeds; k-means pairwise ARI=0.997; GMM pairwise ARI=0.751. |

### Severity-deconfounded residual profiles

Each robust-scaled motor domain was regressed on total motor severity, age, disease duration, LEDD when available, sex, medication state, and PD state. Residual domain profiles were refit with the selected BGMM family. Raw-versus-residual agreement was summarized using ARI and NMI. Severe-tremor and severe-axial patients were matched using nearest neighbors over standardized severity, age, and disease duration.

To address part-whole coupling, sensitivity analyses replaced total motor severity with first principal-component motor burden, leave-one-domain-out total severity, total non-target motor burden, and clinical/covariate-only burden. Each residual profile was refit with the BGMM family, and active component count, ARI, NMI, and domain-wise residualization *R*^2^ were recorded.

### Refit-stability posterior calibration

Patient-blocked bootstrap calibration used 250 refits. Patients were sampled with replacement, all their visits were included, and the selected BGMM configuration was refit to each bootstrap sample. Predicted labels for the full visit table were aligned to original labels using the Hungarian algorithm over the confusion matrix. Empirical consistency was the proportion of refits preserving the original aligned label. Calibration bins compared mean max posterior confidence with mean bootstrap consistency, and was computed as the weighted absolute difference between confidence and consistency. Because states are latent, this is an operational reliability calibration against refit stability, not calibration to biological ground-truth labels.

**Table 3:** Claims, falsifiers, and current evidence. The framework is presented as a set of falsifiable claims rather than as a fixed subtype ontology.

| Claim | Falsifier | Current evidence |
| --- | --- | --- |
| Patient-level state families are stable enough to summarize dominant presentation | Low modal-family fraction | Mean modal-family fraction 0.925 in the full aligned table. |
| States remain dynamic over follow-up | No observed transitions | 25.5% ever transitioned in the full aligned table; PD-only sensitivity subsets show higher transition fractions. |
| States are not only severity strata | Residual model collapses | Residual BGMM retained five observed components across five severity-axis definitions; total-severity raw-vs-residual ARI=0.228. |
| Nominal posteriors are overconfident relative to refit stability | Posterior confidence equals bootstrap consistency | Mean confidence 0.989 versus bootstrap consistency 0.849; =0.140. |
| Uncertainty has clinical information | Entropy/gap not predictive | Entropy/gap-only transition models retained AUC up to 0.765; posterior entropy was associated with Cox transition hazard. |
| Imaging recovers soft posterior signal | JSD poor or ECE high | Fused DaTSCAN+MRI RF JSD=0.209 (95% CI 0.199 - 0.220); severity/demographic+imaging JSD=0.188. |
| External transfer has clinical face validity | No endpoint gradients | BioFIND endpoint gradients reproduced severity, axial, H&Y, and motor-domain gradients after assignment, with bootstrap CIs. |

### Longitudinal future-state prediction

For patients with at least two visits, each visit except the last was used to predict the next visit. Predictors included current motor domains, current axial score, severity, tremor-minus-axial contrast, posterior entropy, posterior gap, max posterior, posterior state probabilities, visit time gap, age, disease duration, LEDD, and current state one-hot indicators. Patient-held-out GroupKFold cross-validation used five folds. Transition baselines included logistic regression, random forest, XGBoost, and LightGBM over feature sets spanning current state only, severity/covariates only, motor domains only, posterior vector only, entropy/gap only, and all features. Future axial baselines included ridge regression, random forest, XGBoost, and LightGBM. Metrics included *R*^2^, RMSE, MAE, AUC, balanced accuracy, Brier score, ECE, calibration slope, and calibration intercept. Autoregressive baseline performance used current axial score to predict next axial score.

Repeated-measures clinical baselines were fit with patient-clustered generalized estimating equations: a binomial exchangeable-correlation GEE for future state transition and a Gaussian exchangeable-correlation GEE for next axial burden. A Cox proportional-hazards model used visit-gap time as duration and next-state transition as the event, with severity, posterior entropy, posterior gap, age, and disease duration as covariates. A categorical HMM over observed state sequences was fit as a generative trajectory baseline. Visit-gap-stratified transition matrices were estimated by quartiles of inter-visit gap.

**Table 4:**
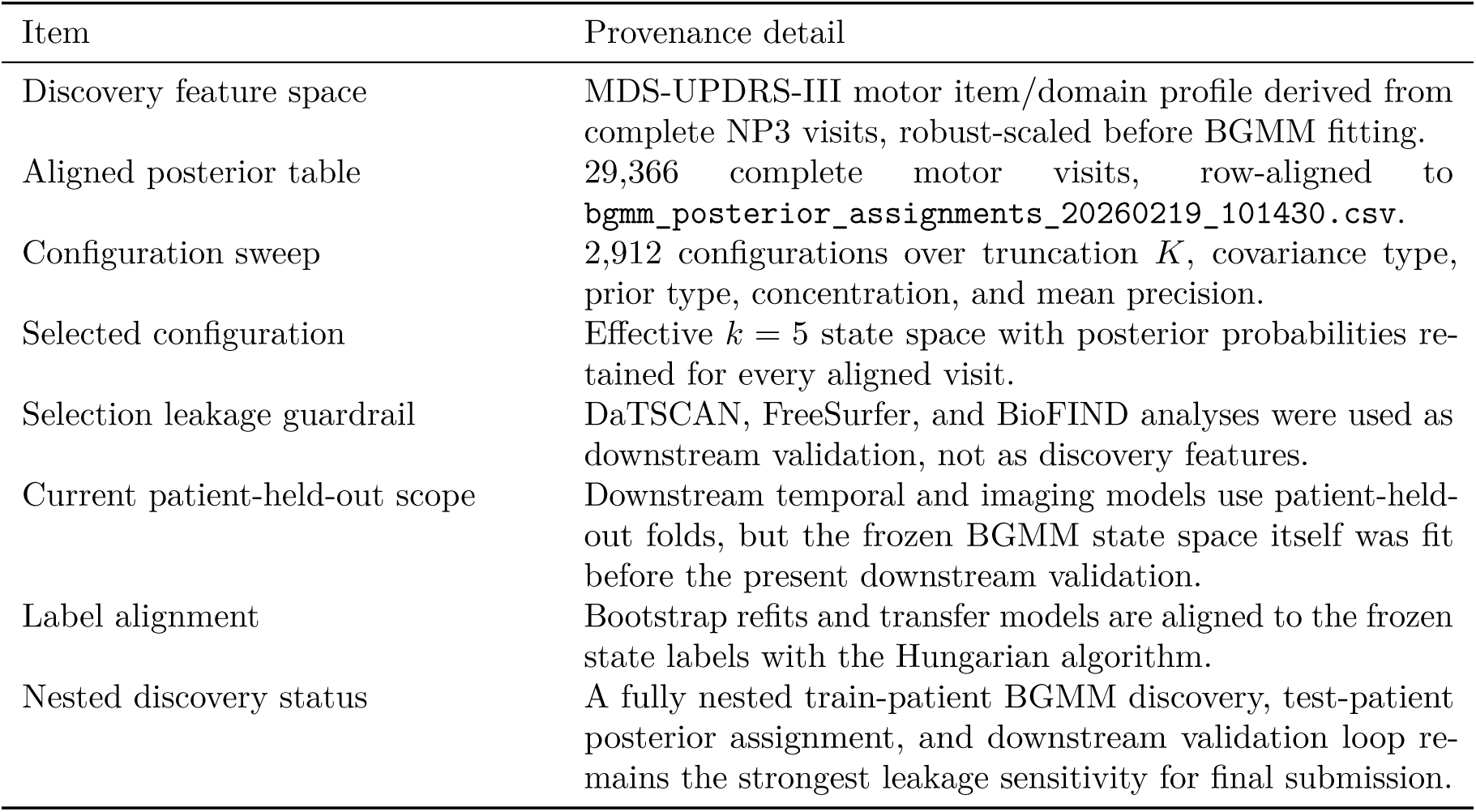
Discovery provenance for the frozen BGMM state space. This table separates the prior discovery step from the downstream patient-held-out and external validation analyses in the current manuscript.

The formal irregular-time extension models transition probabilities as

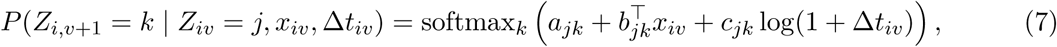

or alternatively as a continuous-time Markov, semi-Markov, or input-output HMM. This extension will report transition intensities and expected dwell time in months or years rather than only visit-indexed transition probabilities.

### Imaging feature construction

DaTSCAN features included bilateral caudate and putamen SBR, anterior putamen SBR when available, mean caudate and putamen SBR, total SBR, caudate/putamen ratio, signed asymmetry indices, and absolute asymmetry indices. FreeSurfer MRI features included eTIV-normalized subcortical volumes from ASEG regions of interest, including putamen, caudate, pallidum, hippocampus, amygdala, thalamus, brainstem, ventricles, ventral diencephalon, and accumbens area.

### Imaging-to-posterior models

Patient-level motor posterior vectors were used as soft labels. RF, XGBoost, and LightGBM regressors were trained under patient-held-out K-fold cross-validation to predict the five-dimensional posterior vector from DaTSCAN features, MRI features, or fused DaTSCAN+MRI features. Predictions were clipped and normalized to the simplex. Evaluation metrics included KL divergence, Jensen–Shannon divergence, Brier score, soft-label ECE, and macro-AUROC for the hard posterior argmax. Feature-group ablations tested DaTSCAN-only, MRI-only, mean-SBR-only, asymmetry-only, severity/demographic-plus-imaging, and imaging residualized for severity. Feature attribution used held-out permutation importance for the fused RF model and SHAP summaries on a 500-patient sample.

### Effect-size-first imaging validation

Patient-level imaging endpoints were modeled with covariate-only and covariate-plus-state linear regressions. Covariates included severity, age, and disease duration. Δ*R*^2^ measured incremental state-associated signal. Kruskal-Wallis *η*^2^ summarized nonparametric effect size across states. Cohen’s *d* was computed for selected severe-axial versus mild-axial contrasts.

### BioFIND transfer and endpoint validation

A transfer BGMM was fit on PPMI NP3 score columns after median imputation and robust scaling. Raw BGMM component labels were mapped to original posterior-state labels with Hungarian alignment. BioFIND motor files were harmonized to PPMI NP3 columns, including a known PN3RIGRL to NP3RIGRL alias. BioFIND state posteriors were predicted, patient-level modal assignments were computed, and endpoint gradients were tested across assigned states. Distribution shift was summarized with Jensen–Shannon divergence between PPMI and BioFIND patient-state distributions. Endpoint associations were summarized using Kruskal-Wallis *η*^2^, FDR-adjusted p-values, and severity-adjusted Δ*R*^2^ when covariates were available. Patient-bootstrap confidence intervals with 1,000 replicates were computed for BioFIND endpoint effect sizes.

### Multimodal latent mixture ablation

BGMMs were fit to five views: motor-only domains, DaTSCAN-only features, MRI-only features, inner-join motor+DaTSCAN+MRI features, and missing-aware motor+DaTSCAN+MRI features with missingness indicators. Agreement with the motor posterior state label was measured using ARI and NMI. Silhouette, active component count, mean max posterior, and latent entropy were reported.

### Baseline stability

K-means, standard GMM, and BGMM baselines were run with 75 random seeds each on patient-level motor-domain features. Agglomerative clustering was run once. For stochastic methods, pairwise ARI and NMI across seeds were reported, along with mean agreement to original posterior-state labels.

### Runtime and software

The professor-review expansion pipeline ran with Python 3.11.8, 120 joblib workers, a 96 tree-model worker cap, 250 patient-blocked BGMM refits, 1,000 bootstrap confidence-interval replicates, five cross-validation folds, 500 estimators for tree/boosting models, and 75 baseline seeds per method on a 144-core ARM Neoverse-V2 node. BLAS thread counts were set to 1 to avoid nested oversubscription. The verified full run completed in 514.3 seconds, with logs saved under logs/prof_review_expansion_20260529_122511.log.

## Data availability

PPMI and BioFIND data are governed by their respective data-access terms and should be requested through the official study portals. Derived aggregate outputs, logs, figures, and manuscript artifacts are stored in results, figures, and logs. Patient-level derived tables may require access-controlled handling before public release.

## Code availability

Code, pipeline, and execution details will be provided upon reasonable request.

## Ethics statement

This retrospective analysis uses de-identified PPMI and BioFIND data. Original cohort studies obtained participant consent and local institutional approvals. The present secondary analysis did not involve participant contact, intervention, or return of individual-level results.

## Acknowledgements

The authors thank the participants, families, investigators, coordinators, imaging cores, data-management teams, and study sponsors of PPMI and BioFIND. The authors also acknowledge the Computing resources were provided by the Texas Advanced Computing Center (TACC) at The University of Texas at Austin.

The authors additionally thank the Peter O’ Donnell Foundation, Jim Holland - Backcountry, and Michael-Connie Rasor.

## Author contributions

All authors contributed equally to this work.

H.M.T. contributed to the computational study design, software implementation, parallel pipeline execution, result verification, figure generation, data analysis, and manuscript preparation. P.Y. contributed to the computational study design, biomedical engineering interpretation, clinical end-point framing, literature synthesis, validation-plan refinement, data interpretation, and manuscript preparation. C.B. contributed to the study conception and design, supervision, computational multimodal clinical data learning strategy, statistical modeling methodology, interpretation of multimodal validation results, and manuscript preparation.

All authors reviewed, revised, and approved the final manuscript.

## Additional information

Supplementary information, source-data tables, reporting-summary material, and code-release paths are organized in the extended data roadmap below.

A shorter version of this work has been accepted for publication in the Proceedings of the International Conference on Medical Image Computing and Computer-Assisted Intervention (MICCAI 2026).

## Extended Data and Supplementary Table Roadmap

**Table 5:** Planned extended data items.

| Item | Content |
| --- | --- |
| Extended Data Fig. 1 | Framework schematic for posterior-calibrated multimodal motor state modeling. |
| Extended Data Fig. 2 | Full patient-level state transition matrix and transition counts. |
| Extended Data Fig. 3 | Posterior calibration bins and entropy/gap distributions. |
| Extended Data Fig. 4 | Imaging-to-posterior feature attribution by modality. |
| Extended Data Fig. 5 | BioFIND endpoint gradients and distribution-shift plot. |
| Supplementary Table 1 | Cohort inclusion audit and candidate PD-only filters. |
| Supplementary Table 2 | Available cohort-sensitivity metrics plus required subgroup rerun metrics for calibration ECE and fused imaging JSD. |
| Supplementary Table 3 | Motor-domain item definitions, residualization coefficients, and leave-one-domain-out severity sensitivity. |
| Supplementary Table 4 | Longitudinal prediction metrics, stronger tabular baselines, and time-scaled transition model. |
| Supplementary Table 5 | Imaging effect-size validation table with bootstrap confidence intervals. |
| Supplementary Table 6 | Imaging-to-posterior permutation importance, feature-group ablation, and per-state calibration. |
| Supplementary Table 7 | Multimodal latent mixture ablation. |
| Supplementary Table 8 | Baseline seed stability benchmark. |
| Supplementary Table 9 | TRIPOD+AI/PROBAST+AI reporting checklist for prediction analyses. |

